# Resolvin E1 derived from eicosapentaenoic acid prevents hyperinsulinemia and hyperglycemia in a host genetic manner

**DOI:** 10.1101/848093

**Authors:** Anandita Pal, Abrar E. Al-Shaer, William Guesdon, Maria J. Torres, Michael Armstrong, Kevin Quinn, Traci Davis, Nichole Reisdorph, P. Darrell Neufer, Espen E. Spangenburg, Ian Carroll, Richard P. Bazinet, Ganesh V. Halade, Joan Clària, Saame Raza Shaikh

## Abstract

**Objective:** Eicosapentaenoic acid (EPA) has recently garnered strong attention given the success of the REDUCE-IT trial, which overturned previous conclusions on EPA and led to its FDA approval for lowering cardiovascular disease risk. Therefore, there is a need to study EPA for cardiometabolic risk factors. Here we focused on EPA’s preventative role on hyperglycemia and hyperinsulinemia.

**Methods:** C57BL/6J male mice were fed a high fat diet in the absence or presence of pure EPA. Mass spectrometry was used to identify how EPA prevents hyperinsulinemia and hyperglycemia that drove subsequent experiments with resolvin E1 (RvE1) across inbred and outbred models.

**Results:** Administration of EPA to C57BL/6J mice prevented obesity-induced glucose intolerance, hyperinsulinemia, and hyperglycemia. Supporting analyses of National Health and Nutrition Examination Survey data showed fasting glucose levels of obese adults were inversely related to EPA intake in a sex-specific manner. We next investigated how EPA improved murine hyperinsulinemia and hyperglycemia. Mass spectrometry revealed EPA overturned the obesity-driven decrement in the concentration of 18-hydroxyeicosapentaenoic acid (18-HEPE) in white adipose tissue and liver. Treatment of obese mice with RvE1, the immunoresolvant metabolite of 18-HEPE, reversed hyperinsulinemia and hyperglycemia through the G-protein coupled receptor ERV1/ChemR23. RvE1’s effects were not mediated by macrophage enrichment in white adipose tissue. Finally, we determined if the metabolic effects of RvE1 were dependent on host genetics. RvE1’s effects on hyperinsulinemia and hyperglycemia were divergent in diversity outbred mice that model human genetic variation. Secondary SNP analyses further revealed extensive genetic variation in human RvE1- and EPA- metabolizing genes.

**Conclusions:** The data suggest EPA prevents hyperinsulinemia and hyperglycemia through the endogenous bioactive metabolite RvE1 that activates ERV1/ChemR23. Importantly, the studies reveal host genetics are an overlooked but critical factor in the metabolic response to RvE1. These results underscore the need for personalized administration of EPA-derived RvE1 based on genetic/metabolic enzyme profiles.

## 1.0 Introduction

Circulating levels of eicosapentaenoic acid (EPA, 20:5n-3) are generally low in the western population [1]. Therefore, increased intake of EPA and other n-3 polyunsaturated fatty acids (PUFA) is hypothesized to ameliorate a range of risk factors that contribute toward cardiometabolic diseases [2]. Notably, EPA has garnered tremendous attention as the recent REDUCE-IT trial showed EPA ethyl esters (Vascepa^®^) substantially reduced the risk of cardiovascular disease in statin-treated patients with elevated triglycerides [3]. As a consequence, the FDA approved EPA ethyl esters for cardiovascular disease risk reduction in select clinical populations. This overturned the dogma on n-3 PUFAs as having no efficacy for lowering the risk of cardiovascular diseases [4].

The effects of n-3 PUFAs on insulin sensitivity and glucose tolerance remain strongly debated. Several randomized clinical trials have failed to establish the benefits of increased n-3 PUFA intake for treating subjects with insulin resistance, which may be due to a range of factors including poor controls, a focus on treatment rather than prevention, and often a neglect for a diversified human genetic makeup [2,5,6]. Furthermore, a unifying mechanism by which n-3 PUFAs such as EPA prevent insulin resistance remains unclear as many studies have relied on heterogenous mixtures of n-3 PUFAs despite evidence that EPA and its long chain counterpart docosahexaenoic acid (DHA) are not structurally or functionally identical [7,8]. To further complicate matters, many preclinical studies rely on levels of n-3 PUFAs that are not easily achievable in humans [2,9,10]. Finally, very little is known about the therapeutic potential of EPA-derived metabolites, which represent the next generation of n-3 PUFA research. Thus, the overarching goal of this study was to address these limitations and to determine the underlying mechanism of action.

We first studied if pure EPA ethyl esters, modeling human pharmacological intake, prevent obesity-induced metabolic impairments using C57BL/6J mice. Supporting analyses on the association between fasting glucose levels and dietary intake of PUFAs was conducted using data from the National Health and Nutrition Examination Survey (NHANES) [11]. We then conducted mass spectrometry based metabolomic and lipidomic analyses to identify targets of EPA ethyl esters, which led to the study of EPA-derived resolvin E1 (RvE1). RvE1 belongs to a family of PUFA-derived endogenous metabolites known as specialized pro-resolving mediators (SPM) that are potent immunoresolvants [12-17]. We specifically investigated if exogenous treatment of RvE1 could reverse hyperinsulinemia and hyperglycemia through the G-protein coupled receptor ERV1/ChemR23. Given that RvE1 is an immunoresolvant, immune profiling experiments using flow cytometry addressed if RvE1 mitigates hyperglycemia via an improvement in adipose tissue inflammation in obesity. Furthermore, we determined if the metabolic effects of RvE1 were dependent on the host genome, which is critical to investigate as humans are genetically heterogenous. To do so, we employed diversity outbred (DO) mice, which are a unique mouse population that model human genetic diversity [18]. In parallel, we conducted supporting SNP analyses by mining the Ensembl database to identify genetic variation of EPA and RvE1 metabolizing genes in humans.

## 2.0 Materials and Methods

### 2.1 Animal models, diets, and RvE1 administration

All murine experiments adhered to IACUC guidelines established by The University of North Carolina at Chapel Hill and East Carolina University for euthanasia and humane treatment in addition to the NIH Guide for the Care and Use of Laboratory Animals. Euthanasia relied on CO_2_ inhalation followed by cervical dislocation. C57BL/6J male mice of 5-6 weeks of age were fed lean control (10% kcal from lard, Envigo TD.160407) or high fat (60% kcal from lard, Envigo TD.06414) diet in the absence or presence of EPA (Cayman, ≥93%) ethyl esters (Envigo TD.160232) for 15 weeks. EPA in the high fat diet accounted for 2% of total energy. The schematic illustrating the generation of the ERV1/ChemR23 mutant allele by CRISPR/Cas9-mediated genome editing is provided in the Supplemental Materials, as described below. These mice and wild type littermate controls were also fed diets from Envigo.

For select studies, C57BL/6J male mice were purchased obese from Jackson at 18 weeks of age. They were acclimatized by feeding lean (10% lard, D12450B) or high fat (60% lard, D12492) diets (Research Diets) for an additional 2-3 weeks prior to conducting experiments. DO male mice (Jackson) from generation 33 were obtained at 4 weeks of age and acclimated for 2 weeks. The DO population is derived from 144 Collaborative Cross lines obtained from Oak Ridge National Laboratory at generations F4-F12 of inbreeding [18].

### 2.2 Body mass and insulin/glucose measurements

Metabolic studies including Echo-Magnetic Resonance Imaging (MRI) experiments were conducted as previously described [19]. Briefly, mice were fasted for 5 hours prior to the establishment of baseline glucose values with a glucometer. For the glucose tolerance test, 2.5g of dextrose (Sigma-Aldrich) per kg lean mass was administered intraperitoneally.

### 2.3 Studies with diversity outbred (DO) mice

Since every DO mouse is genetically unique, each mouse served as its own control. Baseline fasting insulin/glucose measurements were recorded once each DO mouse achieved ∼14 grams of fat mass as measured by Echo-MRI. The mice were then allowed one week to recover and subsequently i.p. injected for 4 consecutive days with 300 ng RvE1 per day [20]. Fasting glucose and fasting insulin were again measured after RvE1 administration.

### 2.4 Untargeted mass spectrometry-based metabolomics

Adipose tissue and liver were homogenized using a bead homogenizer and prepared for metabolomics using previously described methods [21]. Samples were analyzed using liquid chromatography mass spectrometry (LC/MS) and raw data were extracted and processed using Agilent Technologies Mass Hunter Profinder Version B.08.00 (Profinder) software in combination with Agilent Technologies Mass Profiler Professional Version 14 (MPP) as previously described [21-23]. An in-house database containing METLIN, Lipid Maps, Kyoto Encyclopedia of Genes and Genomes (KEGG), and Human Metabolomics Database (HMDB) was used to annotate metabolites based on exact mass, isotope ratios, and isotopic distribution with a mass error cutoff of 10 ppm. This corresponds to annotation at Metabolomics Standards Initiative (MSI) level 3 [24].

To visualize clustering between the dietary groups we ran a principal component analysis (PCA) using all metabolites. We then determined statistically significant metabolites between obese mice and obese mice supplemented with EPA. One of the samples from the high fat diet (HF_105) was an outlier from all the other samples and was excluded from analyses. We then calculated fold changes (EPA/high fat). Next, using the validated significant metabolites with Log2 fold changes ±1.5 we standardized the abundances of the metabolites by assigning a Z-score for each sample based on the distribution of the given metabolite. We utilized the Z-scores to generate heatmaps annotated with the classification of each metabolite.

### 2.5 Targeted mass spectrometry-based metabololipidomics

Analyses of PUFA-derived metabolites of visceral white adipose tissue, liver, and heart was conducted as previously described [25]. Lipid mediators were extracted using Strata-X 33-μm 30 mg/1 ml SPE columns (Phenomenex, Torrance, CA). Quantitation of lipid mediators was performed using two-dimensional reverse phase HPLC tandem mass spectrometry (liquid chromatography–tandem mass spectrometry). All standards and internal standards used for the LC/MS/MS analysis were purchased from Cayman Chemical (Ann Arbor, Michigan, USA). All solvents and extraction solvents were HPLC grade or better.

### 2.6 Flow cytometry analyses of differing immune cell populations in adipose tissue

Epididymal visceral adipose tissue was mechanically chopped and then digested with collagenase Type 4, DNAse 1, and 0.5% fatty acid free bovine serum albumin in phosphate buffered saline (PBS) for 50 minutes at 37°C in an incubator-shaker. Next, 0.5M EDTA was added with RPMI media and the cells were passed through a 70-μm cell strainer rinsed with 2% fetal bovine serum (FBS) in PBS. The cells were centrifuged at 1400 RPM for 5 minutes at 4°C, primary adipocytes were aspirated off from the top layer of the supernatant and the pellet containing the stromal vascular cells (SVC) was then incubated with red blood cell lysis buffer for 1 minute on ice. Two percent FBS-PBS was added to the SVCs and centrifuged at 1400 RPM for 5 minutes at 4°C and then passed through a 40-μm cell strainer. SVCs were stained with the following fluorophore-tagged antibodies obtained from BioLegend: Zombie Aqua, CD45 (PerCP-Cy5.5), CD11b (FITC), CD19 (APC-Cy7), F4/80 (PE), and MHCII (BV421). The following SVC subsets were analyzed using a BD LSRII flow cytometer: CD45^+^CD11b^-^CD19^+^ (B cells), CD45^+^CD11b^+^F4/80^+^MHCII^+^(macrophages),CD45^+^CD11b^+^F4/80^+^MHCII^-^ (macrophages). All data were analyzed in FlowJo and gates were drawn from fluorescence minus one controls.

### 2.7 Analyses of NHANES database

The 2013-2014 NHANES database was mined for daily average intake of PUFAs with respect to age, sex, and BMI. We used Rv3.4.4 with the RNHANES package to retrieve the NHANES database. Graphical packages ggpubr and ggplot2 were used to generate all graphs and statistical annotations. The “Dietary Interview - Total Nutrient Intakes” section of the NHANES database was used to retrieve PUFA intake measurements based on a 24-hour dietary recall questionnaire. OGTT 2-hour glucose measurements were retrieved from the “Current Health Status” section, where BMI was retrieved from “Body Measures”. Tertiles of PUFA intake were calculated corresponding to the probability of intake at 33.3%, 67%, and 100% of the range.

Normality and homogeneity of variance were tested with the Shapiro-Wilks test and Bartlett test respectively. The dataset did not satisfy the assumptions of normality and heteroscedasticity therefore, we utilized a Kruskal-Wallis test followed by a Wilcoxon pairwise test to measure significant differences between tertiles of PUFA intake.

### 2.8 SNP analyses

We used the Biomart tool to mine the Ensembl Variation 98 Human Short Variants (GRCh38.p13) database for single nucleotide polymorphisms (SNPs) with minor allele frequencies at or above 5% that are contained within the 1000 genomes project or the NCBI dbSNP archive. We mined SNPs for the genes listed in the Supplemental Methods. We used the ggplot2 package in R v3.4.4 to plot all the minor allele frequencies by each allele for every gene and chromosome. White lines in the graph represent a break/gap in the minor allele frequency (MAF) distribution, for example a gene may contain MAFs ranging from 0.05-1.5 then 2.5-4.5 with a “break” between the 1.5-2.5 gap. The distances between the SNPs of different genes on the chromosomes was determined using the base pair location at the last SNP of the first gene and the first SNP of the second gene. Distances below 500 kilobases were considered as having a higher likelihood for genetic linkage, as described by the HapMap project Haploview tool. Additional details are in the Supplemental Methods.

### 2.9 Statistics

Data were analyzed using Graph Pad Prism Version 7.0. Statistical significance relied on one-way or two-way ANOVAs followed by a post-hoc Tukey HSD test if the data satisfied the assumptions of normality and homogeneity of variance tested by the Shapiro-Wilks test and Bartlett test respectively. Data that failed the assumption of heteroscedasticity were analyzed using a Welch ANOVA followed by a pairwise Welch T-test with a Bonferroni p-value adjustment. Data sets that did not display normal distributions were analyzed with a Wilcoxon pairwise test. Studies as a function of time that passed the assumptions of normality and heteroscedasticity were analyzed with a two-way ANOVA. For clarity, additional description of analyses are also included with each corresponding methods section above. For all analyses, p<0.05 was considered statistically significant.

## 3.0 Results

### 3.1 EPA limits hyperglycemia, hyperinsulinemia, and improves glucose tolerance of obese male C57BL/6J mice

We first determined if dietary administration of EPA could prevent obesity-induced metabolic impairments of obese male mice. The approach relied on pure ethyl esters of EPA and not mixtures of EPA with DHA that can confound the data. Mice consuming a high fat (HF) diet in the absence or presence of EPA had similar increases in total mass and fat mass compared to lean controls (Fig. 1A). Inclusion of EPA in the diet of obese mice prevented the development of obesity-driven glucose intolerance (Fig. 1B), as quantified by the area under the curve (Fig. 1C). Relative to the HF diet, EPA restored the impairment in fasting glucose (Fig. 1D) and improved fasting insulin levels (Fig. 1E). The HOMA-IR score was also lowered with EPA in the diet, relative to the mice consuming the HF diet (Fig 1F).

**Figure 1.**
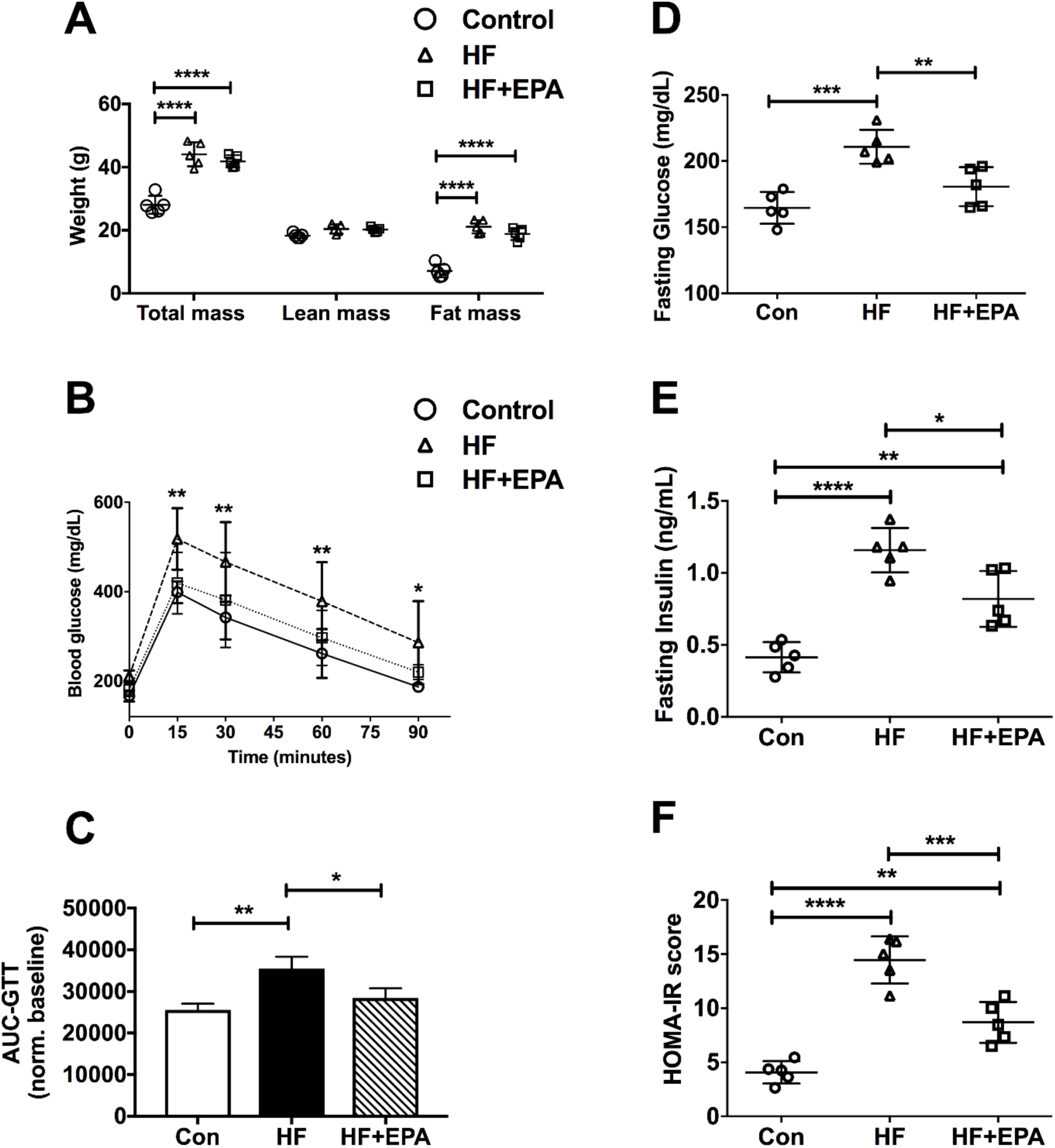
EPA ethyl esters prevent obesity-induced impairments in glucose tolerance, fasting glucose and fasting insulin levels of C57BL/6J mice. (A) Body composition measured by Echo-MRI. (B) Glucose tolerance test performed by intraperitoneal injection of glucose after a 5 hour fast. (C) Area under the curve (AUC), calculated by integration of the curves in B normalized to baseline values. (D) Fasting glucose and (E) fasting insulin levels after a 5 hour fast. (F) HOMA-IR scores. For all measurements, male mice consumed a lean control (Con) diet (O), a high fat (HF) diet (Δ), or a HF diet supplemented with EPA ethyl esters (□). Measurements were conducted at week 13 of intervention. Values are means ± SD. *p<0.05, **p<0.01, ***p<0.001, ****p<0.0001 from one-way ANOVA followed by Tukey’s multiple comparisons test except B, which was a two-way ANOVA followed by a post hoc test.

### 3.2 EPA is associated with improved glucose levels in obese humans in a sex-dependent Manner

To translate the murine data, we analyzed the relation between EPA intake and blood glucose levels during an OGTT in obese humans using data from NHANES. Increased EPA intake was associated with lower glucose levels between the first tertile and the third tertile for obese males (Fig. 2A) but not females (Fig. 2B). Furthermore, we investigated if there was a relationship between DHA and glucose levels. In obese males (Fig. 2C) and females (Fig. 2D), there was no association between DHA and blood glucose levels.

**Figure 2.**
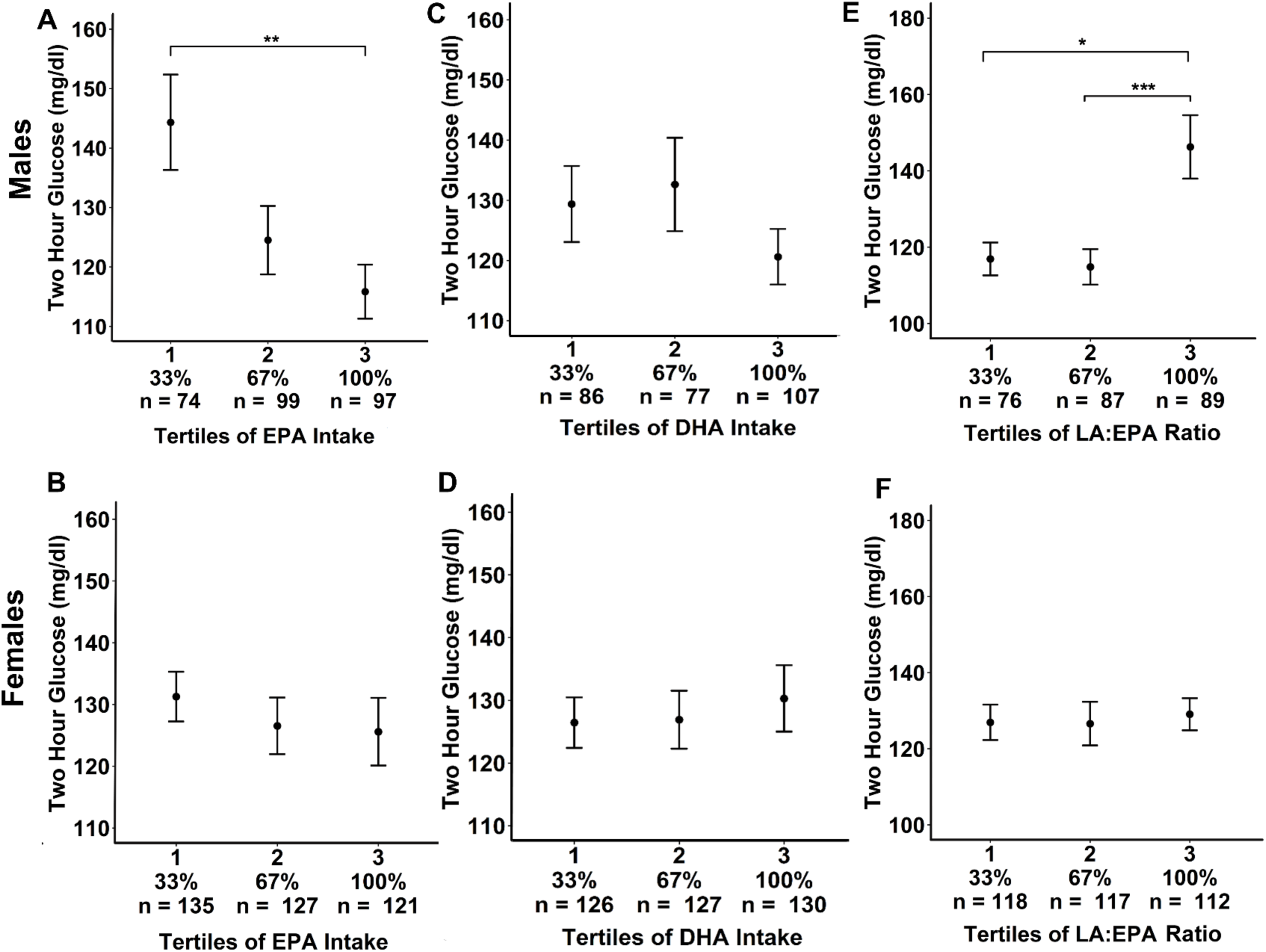
Glucose levels are inversely related to EPA intake in obese men but not women and are dependent on the ratio of LA to EPA. NHANES data on two-hour glucose measurements (mg/dl) from an OGTT were stratified by tertiles of EPA intake in grams for obese (A) males and (B) females. DHA intake is also depicted for (C) males and (D) females. The range of EPA intake for males was 0.0-0.009g for tertile 1, 0.01-0.068g for tertile 2, 0.069g and above for tertile 3. For females, the intake is 0-0.009 for tertile 1, 0.01-1.51g for tertile 2, 1.52g and above for tertile 3. The range of DHA intake for males is 0.01-0.05g for tertile 1, 0.06-1.49g for tertile 2, and 1.5g and above for tertile 3. The range of DHA intake for females was 0.01-0.03g for tertile 1, 0.04-2.35g for tertile 2, and 2.36g and above for tertile 3. Subjects were adults (18 years and older) and had a BMI of 30 and above. Tertiles of the ratio of LA to EPA are presented for obese (E) males and (F) females. The tertiles correspond to 33%, 67%, and 100% of the range of LA to EPA intake for subjects older than 18 and a BMI of 30 and above. Values are means ± SEM *p<0.05, **p<0.01, ***p<0.001. from Wilcoxon pairwise test. Number of subjects for each tertile is listed on the x-axis.

The metabolism of EPA relies on some of the same immune responsive and metabolic enzymes used by n-6 PUFAs such as linoleic acid (LA) [26,27]. Thus, we further mined the NHANES data to determine if there was a relationship between the ratio of LA to EPA on fasting glucose levels. Tertiles of the LA to EPA ratio are plotted for obese men (Fig. 2E) and women (Fig. 2F). The positive association between EPA and glucose levels was diminished and strikingly, at the highest ratio of LA to EPA in men (Fig. 2E), but not women (Fig. 2F), blood glucose levels were increased relative to the first two tertiles.

### 3.3 Mice consuming EPA have a distinct metabolome compared to obese mice in the absence of EPA

As the mechanism of action for EPA ethyl esters is likely to be pleiotropic, we conducted metabolic profiling of mice consuming the experimental diets. PCA plots revealed a clear distinction between the control, HF, and HF + EPA ethyl ester diets for visceral white adipose (Fig. 3A). EPA ethyl esters were predominately incorporated into triglycerides with some uptake into diglycerides and phosphatidylcholine (Fig. 3B, 3C). In the liver, PCA plots also showed a clear distinction between the HF and HF + EPA ethyl ester diets (Fig. 3D). EPA ethyl esters appeared to have a broad effect on the liver metabolome (Fig. 3E, 3F). EPA acyl chains were likely distributed into triglycerides, phosphatidylcholine, phosphatidylethanolamine, and anandamide (Fig. 3F). Overall, these results showed that EPA ethyl esters are incorporated into several lipid pools, which would then differentially influence metabolic pathways, particularly in the adipose tissue and liver. The full metabolite names, p-values, fold changes, and quantifications are in Supplemental Tables 1 and 2.

**Figure 3.**
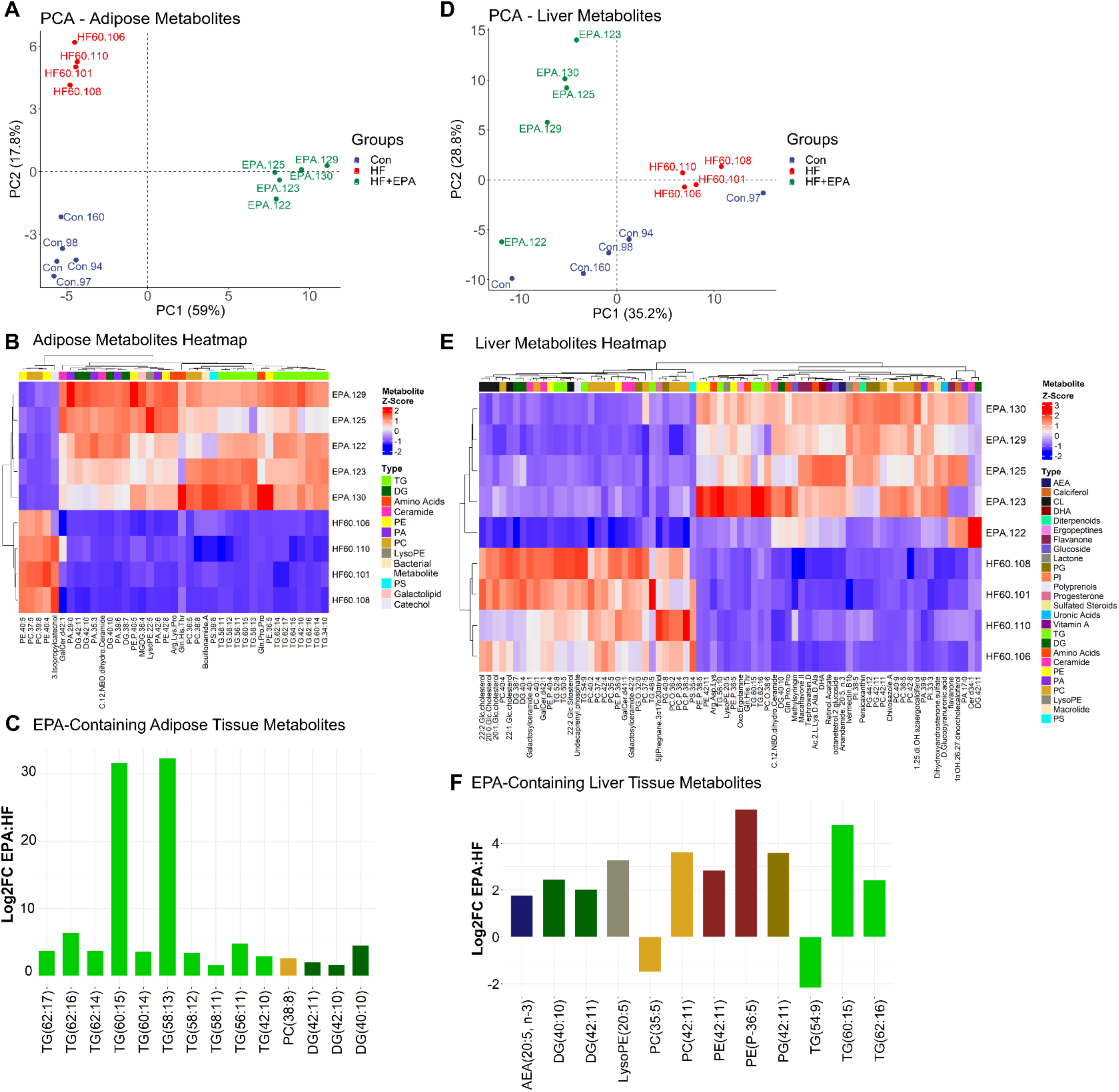
EPA ethyl esters have a distinct metabolomic profile in white adipose tissue and liver of obese C57BL/6J male mice. C57BL/6J male mice consumed a control (Con), high fat (HF) and HF+EPA ethyl ester diet. (A) PCA plot of validated adipose tissue metabolites between control, HF, and HF+EPA samples. (B) Heatmap of Z-scores from significant adipose tissue metabolites with ±1.5 fold-change. (C) Log2 fold change graph of EPA-containing (20:5) adipose metabolites. (D) PCA plot of validated liver metabolites between control, HF, and HF+EPA samples. (E) Heatmap of Z-scores from significant liver metabolites with ±1.5 fold-change. (F) Log2 fold change graph of EPA-containing (20:5) adipose metabolites. Heatmap legends on the right hand side of (B) and (E) show each metabolite’s classifications: triglyceride (TG), diacylglycerol (DG), phosphatidylethanolamine (PE), lysophosphatidylethanolamine (LysoPE), phosphatidic acid (PA), phosphatidylcholine (PC), phosphatidylserine (PS), phosphatidylglycerol (PG), phosphatidylinositol (PI), arachidonoylethanolamine (AEA), cholesterol (CL), and docosahexaenoic acid (DHA). Full metabolite names are provided in the supplemental.

### 3.4 12-HEPE and 18-HEPE levels are lowered with obesity and reversed with EPA

We next conducted targeted metabololipidomic analyses to further study the effects of EPA on the adipose and liver lipidomes. The HF diet and HF diet + EPA modulated several n-3 and n-6 PUFA-derived metabolites in these tissues. The HF diet amplified several white adipose tissue n-6 PUFA derived mediators, many of which are pro-inflammatory, relative to the lean control (Fig. 4A). These included 11(12)-EET, 14(15)-EET, 15R-LXA_4_, 6-*α*-PG, 8-iso-15R-PGF2*α*, 8-iso-PGF2*α*, carboxylic TXA_2_, 8S-HETE and 15S-HETE. Inclusion of EPA in the HF diet also amplified some but not all of these metabolites (Fig. 4A). Markedly, the HF diet lowered the levels of the EPA-derived 18-HEPE, which was reversed with EPA in the diet. 18-HEPE levels were elevated by 5.6 and 32.6-fold relative to the lean and HF diets, respectively (Fig. 4A). EPA also strongly upregulated the concentration of 12-HEPE in white adipose tissue by ∼58-fold.

**Figure 4.**
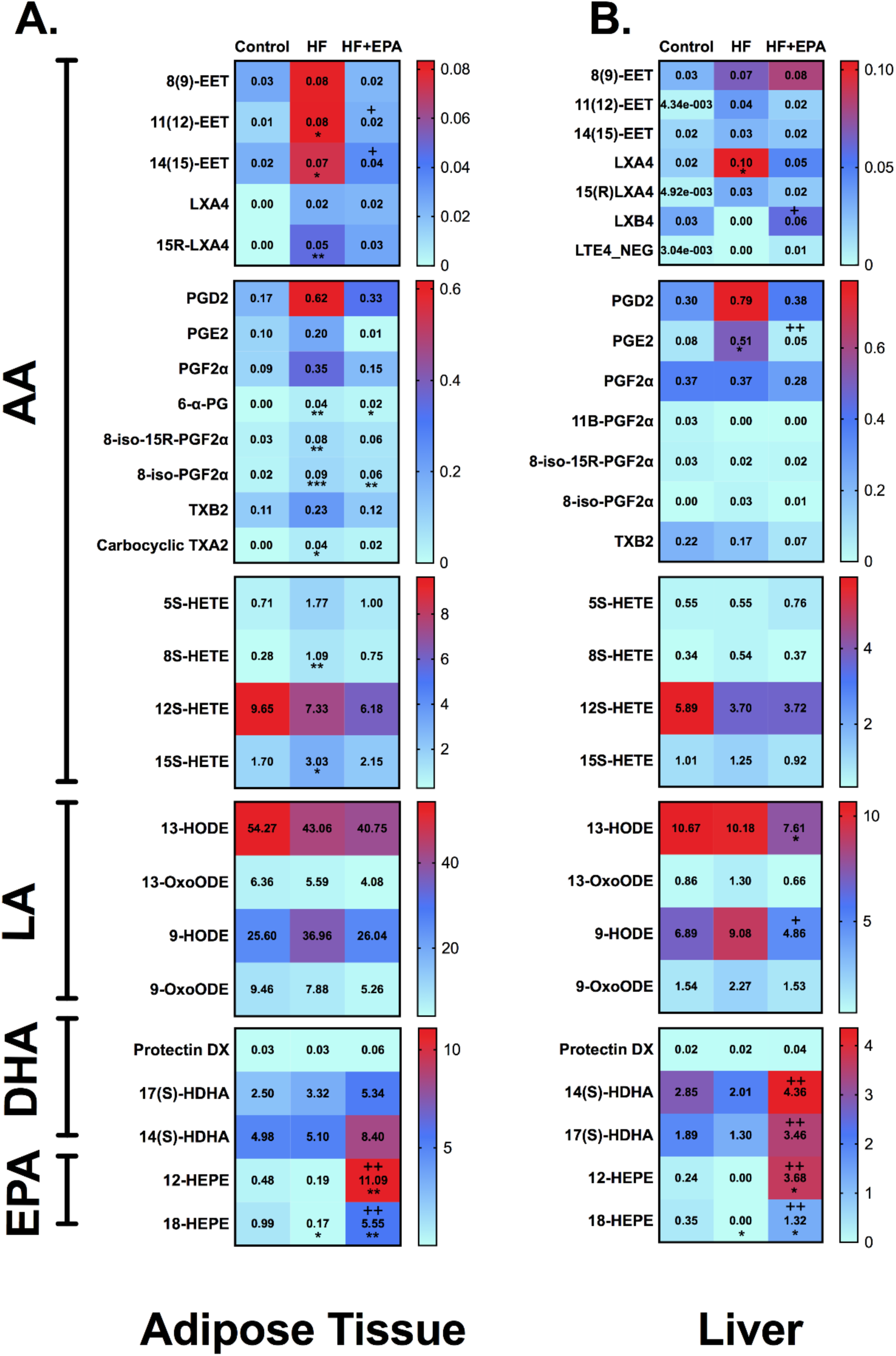
EPA ethyl esters reverse the effects of murine obesity on the concentration of 18-HEPE. Mass spectrometry based metabololipidomic analyses of (A) visceral white adipose tissue and (B) liver. Metabolites from eicosapentaenoic acid (EPA), docosahexaenoic acid (DHA), linoleic acid (LA) and arachidonic acid (AA) are depicted in the heat map. Male C57BL/6J mice consumed experimental diets for 15 weeks. N=4-5 mice per diet. Data are average. *p<0.05, **p<0.01, ***p<0.001 as compared to a control diet and ^+^p<0.05, ^++^p<0.01 as compared to the high fat diet. Statistical analyses for these data are described in the methods section.

In the liver, the n-6 PUFA-derived LXA_4_ and PGE_2_ were increased with the HF diet relative to the lean control (Fig. 4B). Notably, 12-HEPE and 18-HEPE levels were undetectable with the HF diet compared to the control. In contrast, EPA lowered PGE_2_ and 9-HODE levels relative to the HF diet. EPA dramatically increased the levels of 12-HEPE and 18-HEPE compared to the lean and HF diets (Fig. 4B). EPA also elevated the levels of 14-HDHA and 17-HDHA relative to the HF diet.

We also analyzed the heart to determine if the effects of EPA were limited to white adipose tissue and liver. In the heart, EPA had some modest effects on the number of n-6 PUFA derived metabolites modified relative to the HF diet (Suppl. Fig. 1). However, similar to the adipose tissue and liver, EPA increased the concentration of 12-HEPE and 18-HEPE by up to 11-27 fold relative to the lean and HF diets, respectively (Suppl. Fig. 1). Overall, the robust amplification of 18-HEPE levels set the basis for subsequent experiments with RvE1, the downstream and immune responsive bioactive of 18-HEPE.

### 3.5 RvE1 treatment improves hyperinsulinemia and controls hyperglycemia of inbred mice in a manner that is dependent on the receptor ERV1/ChemR23

Given that EPA strongly upregulated 18-HEPE, we subsequently investigated the effects of RvE1 on fasting insulin and glucose using C57BL/6J mice. Furthermore, we tested the hypothesis that the effects of RvE1 could be mediated by one of its two receptors known as ERV1/ChemR23. Therefore, we generated ERV1/ChemR23 knockout (KO) mice and wild type (WT) littermates to measure the effects of RvE1 on hyperinsulinemia and hyperglycemia. Figure 5A and 5B respectively show the experimental scheme and the ERV1/ChemR23 deletion allele. Body composition did not differ between obese WT and ERV1/Chem23 KO mice administered vehicle control or RvE1 (Fig. 5C). RvE1 treatment improved fasting glucose levels of WT but not ERV1/ChemR23 KO mice compared to obese mice (Fig. 5D). We also tested fasting insulin levels in a subset of our WT and KO mice via an ELISA. Fasting insulin levels were improved in response to RvE1 in WT obese mice but not in ERV1/ChemR23 KO obese mice relative to obese mice that received the vehicle control (Fig. 5E).

**Figure 5.**
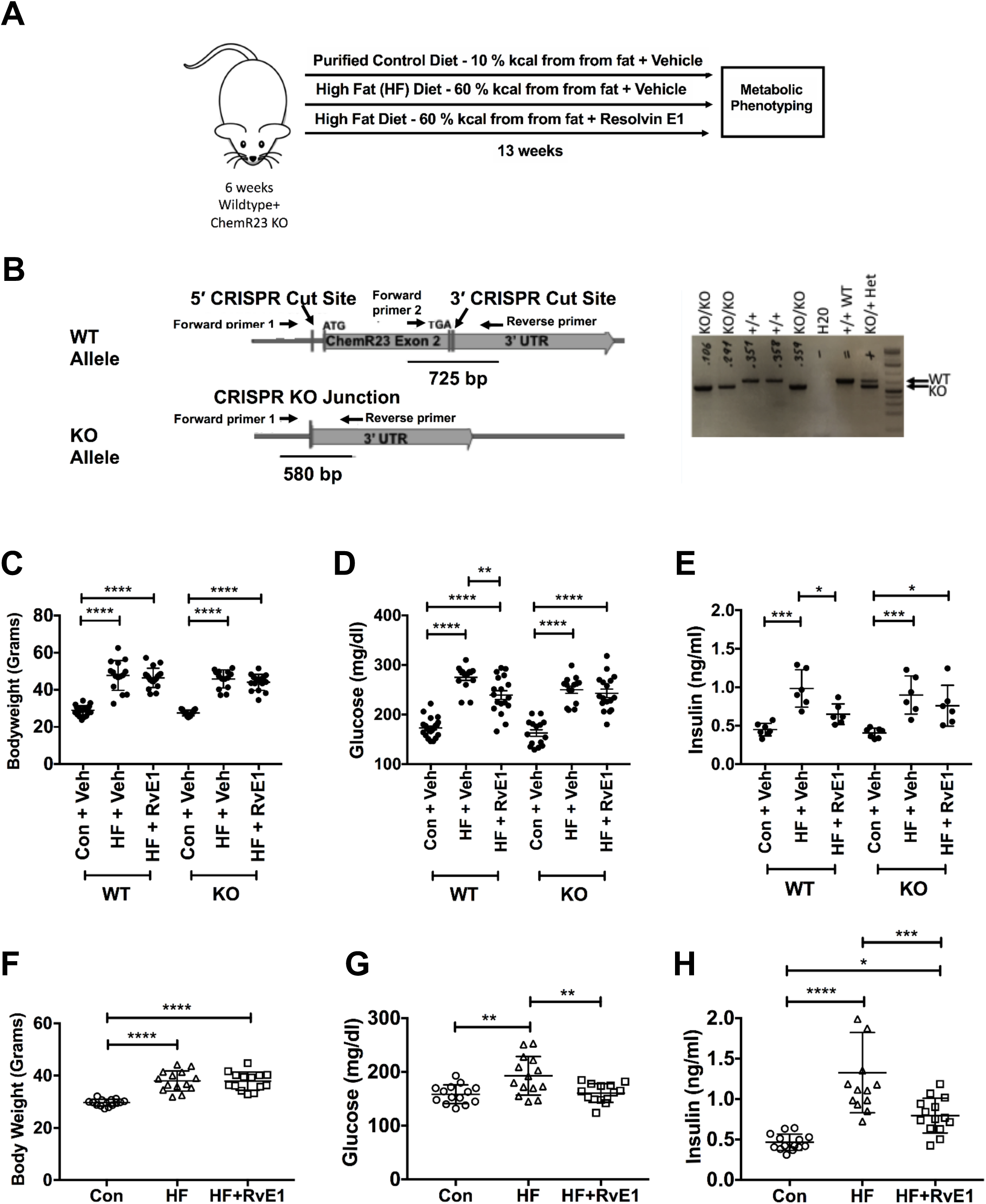
RvE1 treatment improves obesity-driven impairments in fasting glucose and insulin of C57BL/6J mice mediated by the receptor ERV1/ChemR23. (A) Study design with ERV1/ChemR23 knockout mice and their wildtype littermates administered RvE1 or vehicle control after being fed a high fat diet or lean control diet. (B) Illustration of ChemR23 deletion allele and genotyping. (C) Body weight (D) fasting glucose and (E) fasting insulin levels of male wild type (WT) and ERV1/ChemR23 knockout (KO) mice consuming a lean control diet (Con) (O) or high fat (HF) diet in the absence (Δ) or presence (□) of RvE1. Corresponding (F) body weight, (G) fasting glucose and (H) fasting insulin levels of C57BL/6J male mice purchased obese from Jackson Laboratories. All measurements were made at 13-14 weeks of dietary intervention. N=14-18 mice per diet (C,D), N=6-7 mice per diet (E), N=14 mice per diet (F-G). Values are means ± SD. *p<0.05, **p<0.01, ***p<0.001, ****p<0.0001 by one-way ANOVA followed by Tukey’s multiple comparisons test.

We also determined if the RvE1 treatment limits hyperglycemia and hyperinsulinemia using C57BL/6J as a second diet model system (diets from Research Diets) that are prone to obesity and obesity-induced metabolic defects, which were mice obtained obese from Jackson Laboratories. RvE1 treatment of obese mice had no effect on body weight (Fig. 5F) compared to the mice on a high fat diet. RvE1 restored fasting glucose (Fig. 5G) and improved fasting insulin levels relative to obese mice (Fig. 5H).

Mechanistically, immune cells such as B lymphocytes and macrophages in white adipose tissue have a central role in regulating glucose metabolism and thereby metabolic homeostasis. Furthermore, previous work shows that resolvin D1 improves glycemic control by targeting the number of M1- and M2-like macrophages in white adipose tissue [13]. Thus, we investigated if RvE1 was preventing the enrichment of key immune cells in visceral white adipose tissue. The flow cytometry gating strategy is depicted in Fig. 6A and fluorescence minus one controls are shown in Suppl. Fig. 2. For this select study, we only compared mice consuming a HF diet in the absence or presence of RvE1 as significant pooling of lean mice was required. Strikingly, the analyses revealed RvE1 had no effect on the percentage of CD45^+^CD11b^-^CD19^+^ B cells (Fig. 6B), CD45^+^CD11b^+^F4/80^+^MHCII^+^ macrophages (Fig. 6C), or CD45^+^CD11b^+^F4/80^+^MHCII^-^ macrophages (Fig. 6D). Similarly, there was no effect of RvE1 on the number of CD45^+^CD11b^-^CD19^+^ B cells (Fig. 6E), CD45^+^CD11b^+^F4/80^+^MHCII^+^ macrophages (Fig. 7F), or CD45^+^CD11b^+^F4/80^+^MHCII^-^ macrophages (Fig. 6G). We also probed if RvE1 had any impact on select inflammatory and metabolic transcripts. qRT-PCR analysis revealed no effect of RvE1 on the expression of adiponectin, *Tnfα, IL10*, and *GLUT-4* (data not shown).

**Figure 6.**
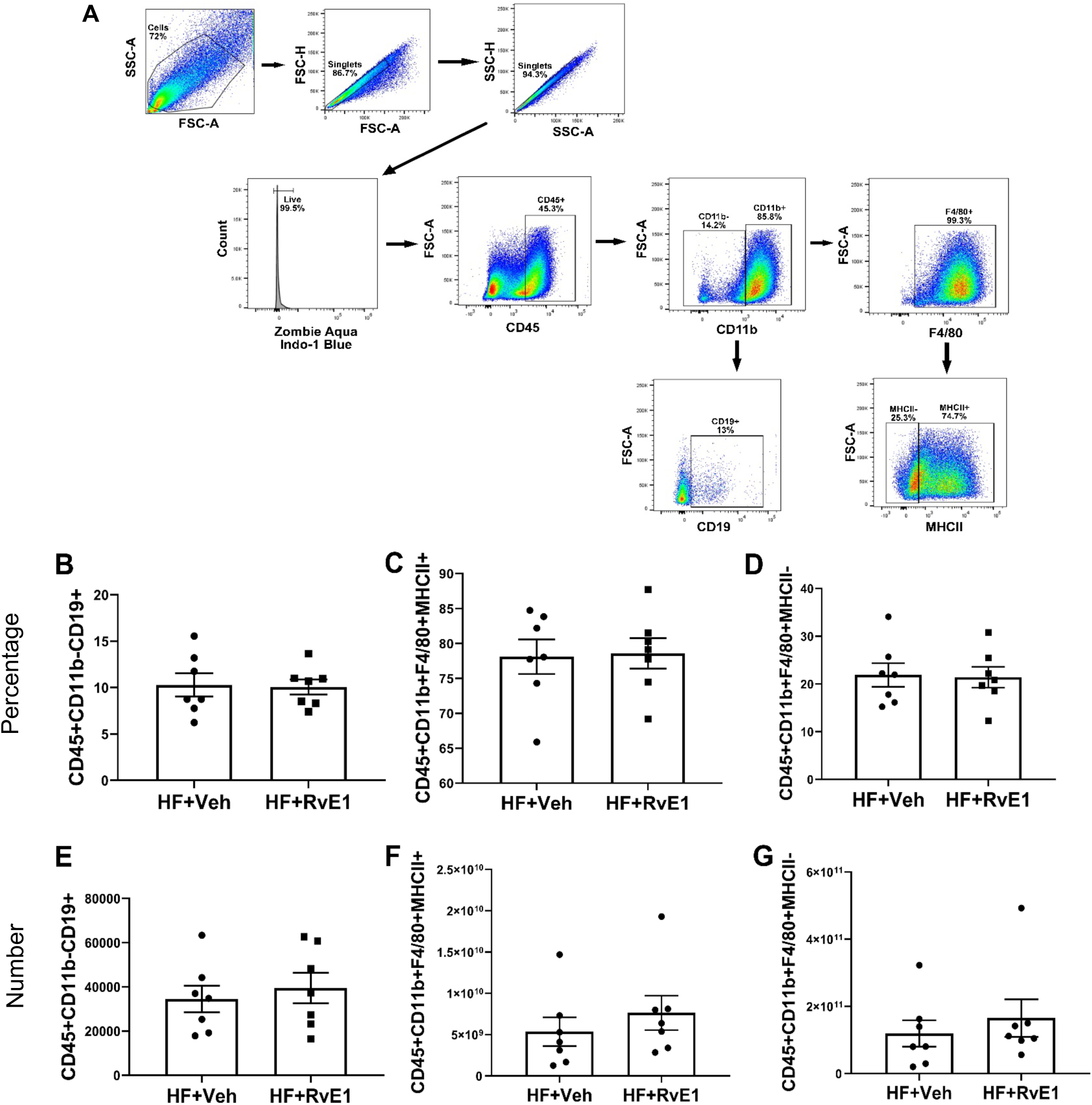
The metabolic improvement with RvE1 is not driven by a reduction in the enrichment of select pro-inflammatory immune cells in white adipose tissue. Flow cytometry gating strategy for B lymphocytes and macrophage populations in the white adipose tissue of obese C57BL/6J male mice purchased from Jackson Laboratories. All measurements were made at 13-14 weeks of dietary intervention. The percentage of (B) CD45^+^CD11b^-^CD19^+^ B cells (C) CD45^+^CD11b^+^F4/80^+^MHCII^+^ macrophages and (D) CD45^+^CD11b^+^F4/80^+^MHCII^-^ macrophages. The number of (E) CD45^+^CD11b^-^CD19^+^ B cells, (F) CD45^+^CD11b^+^F4/80^+^MHCII^+^macrophages and (G) CD45^+^CD11b^+^F4/80^+^MHCII^-^ macrophages. N=6-7 mice per diet. Values are means ± SD.

**Figure 7.**
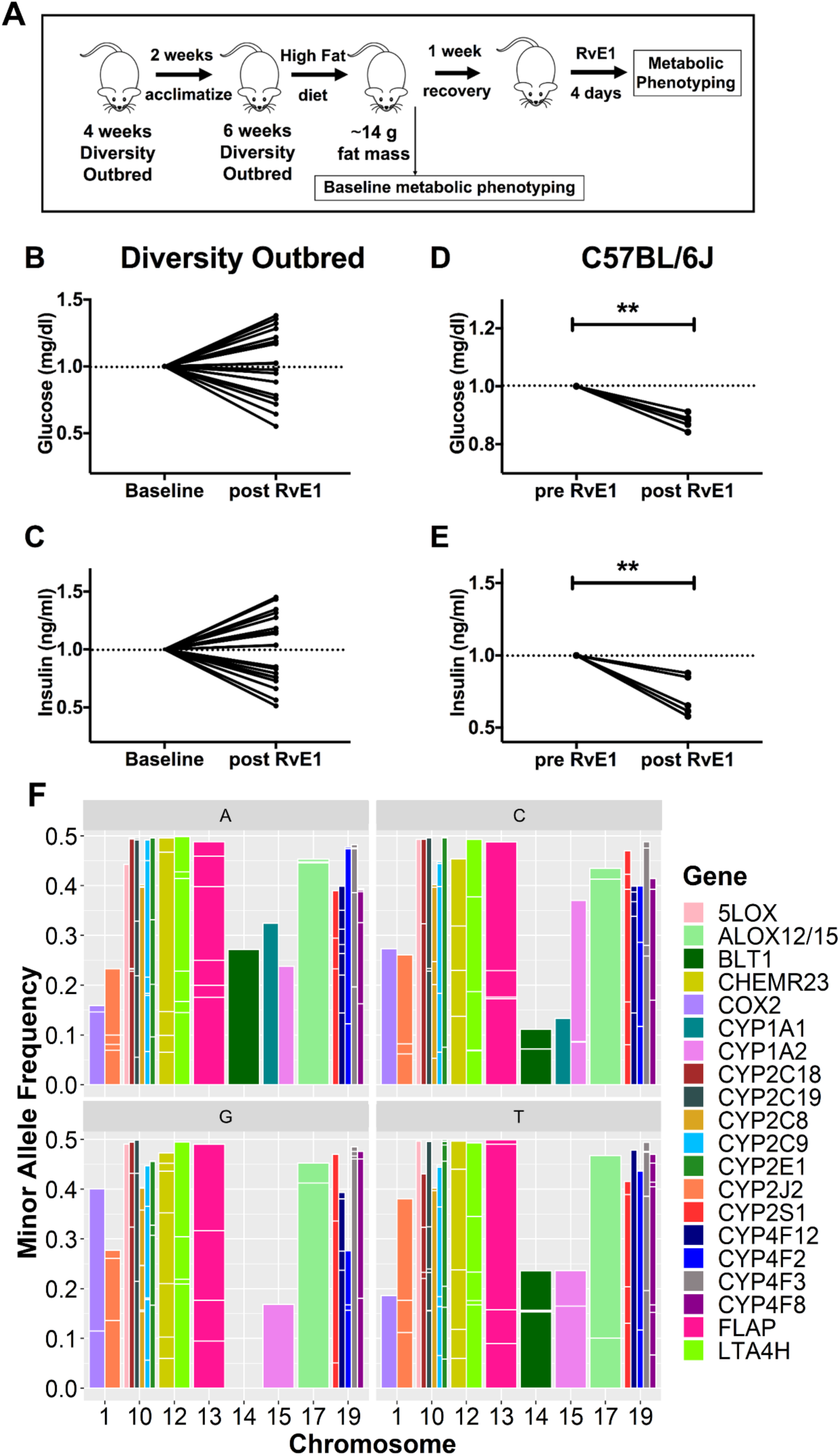
Host genetics have a critical role in the metabolic response to RvE1. (A) Schematic representation of study design with diversity outbred mice. Fasting glucose and insulin measurements were obtained prior to (baseline) and after 4 days of RvE1 administration. (B) Fasting glucose and (C) fasting insulin levels after a 5 hour fast. Corresponding studies with C57BL/6J mice using the same experimental design of intervention are depicted for (D) fasting glucose and (E) fasting insulin. All data are plotted as the fold change in glucose and insulin relative to baseline. N =19 DO mice and N= 6 C57BL/6J mice. **p<0.01 by a paired two-tailed t test. (F) Data were mined from the 1000 genomes and dbSNP human variants databases. Analyses show SNPs in the EPA and RvE1 metabolizing genes stratified by each minor allele (A,C,G,T). The SNPs in each gene are plotted by minor allele frequencies and the chromosome that contains the SNP. The major genes that metabolize EPA and/or RvE1 are depicted by differing colors.

### 3.6 Host genetics define the treatment response to RvE1

Humans are metabolically heterogeneous with a diversified genetic makeup therefore, we determined whether host genetic differences lead to variations in the fasting insulin and fasting glucose response to RvE1. For these studies, we used the translational DO mouse model that mimics human genetic diversity and variability [18]. Similar to the human population, administration of a high fat diet led to large variations in body weight gain of DO mice. Thus, we optimized an experimental design (Fig. 7A) to selectively measure the effects of RvE1 on fasting insulin and fasting glucose. Mice that achieved ∼14g of fat mass (measured via Echo-MRI) over the course of the dietary intervention with a high fat diet were used for these studies. The rationale for selecting ∼14g of fat mass was based on the studies with obese C57BL/6J mice (Figs. 1, 5) that were in this range of fat mass. Body weight gain of the DO mice is depicted in Suppl. Fig. 3. Relative to baseline, RvE1 improved fasting glucose (Fig. 7B) and fasting insulin (Fig. 7C) levels in only half of the obese DO mice. In contrast, using the same experimental design with C57BL/6J mice, fasting glucose (Fig. 7D) and fasting insulin (Fig. 7E) were more uniformly improved in response to RvE1 treatment.

The results with the DO mice led us to further investigate if there is strong genetic variation in RvE1- and EPA-metabolizing genes in humans. We mined the Ensembl database containing the dbSNP archive and 1000 genomes data (Suppl. Table 3). We extracted all the CYP450 enzymes that have the capacity to metabolize EPA, further downstream enzymes leading to the production of E-series resolvins (COX2, ALOX5, FLAP, ALOX12/15, LTA4H), and the two RvE1 receptors (ChemR23 and BLT1) [28-30]. The analyses show a large range of minor allele frequencies where SNPs for each gene are contained in chromosomes 1, 10, 12-15, 17 and 19 with BLT1 lacking SNPs in chromosome 14 (Fig. 7F). Genes with lower ranges of minor allele frequencies (MAF) ranging from 0.05 (5%) – 0.38 (38%) include BLT1, COX2, CYP2J2, CYP1A1 and CYP1A2. Surprisingly, all other genes contained many high MAFs with numerous SNPs in the 0.4 (40%) - 0.5 (50%) range (Fig. 7F). Moreover, the CYP450 genes contained many SNPs in close proximity (<500 kilobases) on the same chromosome, these include: CYP2C18 and CYP2C19 (∼27 Kb apart), CYP2C19 and CYP2C9 (∼84 Kb), CYP2C9 and CYP2C8 (∼47 Kb), CYPA1A1 and CYPA1A2 (∼24 Kb), CYP4F8 and CYP4F3 (∼10 Kb), CYP4F3 & CYP4F12 (∼10 Kb), and CYP4F12 and CYP4F2 (∼181 Kb). Taken together, these results showed high population variance in EPA- and RvE1-metabolizing genes.

## 4.0 Discussion

There is growing evidence from humans and rodents that obesity impairs the biosynthesis of SPMs and their precursors, which are predominately generated from dietary n-3 PUFAs including EPA [14,19,31,32]. The loss of SPMs contributes toward a range of cardiometabolic complications including chronic inflammation, hepatic steatosis, insulin resistance, susceptibility to infection, and delayed wound healing [13,15,25,32-35]. Therefore, there is an unmet need to understand how specific dietary n-3 PUFAs through the biosynthesis of their downstream metabolites regulate outcomes in cardiometabolic and inflammatory diseases.

This study advances the field by demonstrating that administration of EPA ethyl esters can prevent hyperinsulinemia and control hyperglycemia through the actions of RvE1 in a host genetic dependent manner. One notable outcome were the data from the NHANES analyses, which underscore the need for preventative precision nutrition studies with EPA. Longitudinal studies with EPA that are focused on prevention, prior to the onset of insulin resistance, are limited. One pilot study demonstrated that administration of n-3 PUFAs to healthy human volunteers prevented insulin resistance induced with a glucocorticoid [36]. The NHANES analyses revealed those subjects with lowest consumption of EPA, but not DHA, had the highest glucose levels in a sex-specific manner. The positive effects of EPA were mitigated when the intake of LA was high, which is common in the western diet. While these data are associative, they point to the importance of discriminating EPA from DHA and accounting for LA levels in clinical studies and trials. LA biosynthesis and metabolism requires some of the same enzymes used for EPA synthesis and metabolism, including the endogenous biosynthesis of downstream metabolites such as RvE1 [26,27]. Moreover, LA’s consumption in the western diet is 14-18 times the amount required to prevent LA deficiency [2,37] and LA’s effects on insulin sensitivity are debated [38]. LA is an essential signaling PUFA for human and animal health but its overconsumption may contribute toward chronic inflammation and metabolic impairments.

The data with EPA ethyl esters challenge previous findings. Earlier studies reported a reduction in fat mass with EPA-containing oils, which we did not observe [39,40]. The differences in results using EPA ethyl esters compared to previous findings on body weight maybe due to the concentration/purity of EPA, duration, and use of EPA/DHA mixtures. EPA driving an upregulation in the concentration of RvE1’s precursor is highly consistent with a recent clinical trial to show that consumption of an n-3 PUFA supplement promoted a strong upregulation of SPMs within 24 hours [41].

A second major advancement are the data to show that RvE1 could improve hyperinsulinemia and hyperglycemia via ERV1/ChemR23. RvE1 treatment is likely exerting its effects through multiple mechanisms. There is evidence that RvE1 blocks signaling through BLT1, the receptor for the arachidonic acid-derived LTB4 [16]. The pro-inflammatory mediator LTB4 serves as a chemokine for select immune cell populations that exacerbate glucose intolerance, particularly in white adipose tissue [42]. We did not find evidence that RvE1 is targeting the enrichment of immune cell populations in the adipose tissue as reported for resolvin D1 [13]. The data are consistent with a study to show that protectin DX could improve insulin resistance independent of an effect on adipose tissue inflammation [17]. We did find that RvE1’s parent compound EPA lowered some pro-inflammatory n-6 PUFA-derived mediators, which could also influence the long-term inflammatory milieu.

RvE1 activation of ERV1/ChemR23 may be further inhibiting signaling through chemerin, an adipokine that binds ERV1/ChemR23 [43]. At the organ and cellular level, RvE1 can improve inflammation though the targeting of intestinal alkaline phosphatase and specific cell types including neutrophils and through the production of further downstream metabolites such as 18-oxo-RvE1 [32,35,44,45]. The effects of RvE1 are likely to occur simultaneously with other EPA-derived metabolites. Notably, we observed EPA strongly upregulated the concentration of 12-HEPE, which was recently identified to improve glucose metabolism [46].

There is some ambiguity among studies regarding the effects of RvE1 on hyperglycemia. Overexpression of ERV1/ChemR23 in myeloid cells improved hyperglycemia and hepatic steatosis of male mice [47]. However, RvE1 administration to wild type mice at a dose of 2ng/g body weight twice weekly for four weeks did not improve hyperglycemia [47]. Our studies relied on a dose of RvE1 (300ng/mouse) for four consecutive days, which may explain the positive effects. Overall, our data on RvE1 were in agreement with the literature to show that select DHA-derived SPMs improve insulin resistance [13,17,33,48,49, 50]. For instance, the DHA-derived SPM protectin DX and resolvin D1 enhances glucose tolerance [13,17].

A final advancement from this study is that DO mice, which model human genetic diversity, respond in a divergent manner upon RvE1 administration. The results open a new area of investigation by suggesting that RvE1 is unlikely to have a uniform positive effect in all obese humans. These results underscore the need for precision treatment in the human population. Overall, little is known about the role of host genetics on SPM biology. One study highlights the importance of genetic variation by demonstrating that obese subjects with a C allele in the rs1878022 polymorphism of ERV1/ChemR23 receptor confers protection from adipose tissue inflammation [51].

The study with DO mice directly informs emerging clinical trials with EPA-derived metabolites on the need to account for the host genome. To exemplify, a recent placebo controlled randomized trial with a 15-hydroxy EPA ethyl ester (Epeleuton) showed an improvement in glycemic control, HbA1c, and inflammation in adults with non-alcoholic fatty liver disease [52]. Therefore, studies such as this one may find more robust effects in future precision clinical trials. To support this conclusion, mining the 1000 genomes and dbSNP databases revealed a large range of minor allele frequencies in the EPA and RvE1-metabolizing genes. Most of the genes analyzed reach minor allele frequencies close to 50%, indicating large population variance in the EPA-RvE1 pathway. Furthermore, close proximity of the CYP450 enzyme SNPs suggest potential genetic linkage in many of the CYP450 variants that can potentially influence metabolism of EPA and its downstream metabolites. The results provide strong groundwork for future genetics studies that will establish candidate genes regulating the metabolic response to RvE1. This will allow investigators to establish ‘responders’ from ‘non-responders’ to EPA-derived metabolites in humans.

## 5.0 Conclusions

In summary, the results provide strong evidence that the EPA-RvE1 axis has a critical role in controlling insulin and glucose homeostasis, which may be a preventative target for cardiometabolic diseases. The results across model systems highlight the need for future prevention and treatment studies that account for the role of host genetics in the metabolism of RvE1 and other EPA-derived metabolites.

## Author contributions

A.P. investigation, formal analysis, visualization, validation, data curation, writing of original draft; A.E.A. investigation, formal analysis, visualization, validation, data curation, writing parts of original draft, W.G. investigation; M.T. investigation, validation; M.A. investigation; K.Q. investigation, validation; T.D. investigation; N.R. methodology, project administration, funding acquisition; P.D.N. resources, funding acquisition; E.E.S. resources, funding acquisition; I.C. investigation, visualization; R.B. writing-review and editing; G.V.H. writing-review and editing, funding acquisition; J. C. writing-review and editing;, and S.R.S. methodology, formal analysis, writing-review and editing, supervision, project administration, funding acquisition.

## Acknowledgements

We thank Dr. Dale Cowley from the UNC Animal Models Core for his assistance in generating the ERV1/ChemR23 KO mice.

