## Supplemental Figures for "Resolvin E1 derived from eicosapentaenoic acid prevents hyperinsulinemia and hyperglycemia in a host genetic manner"

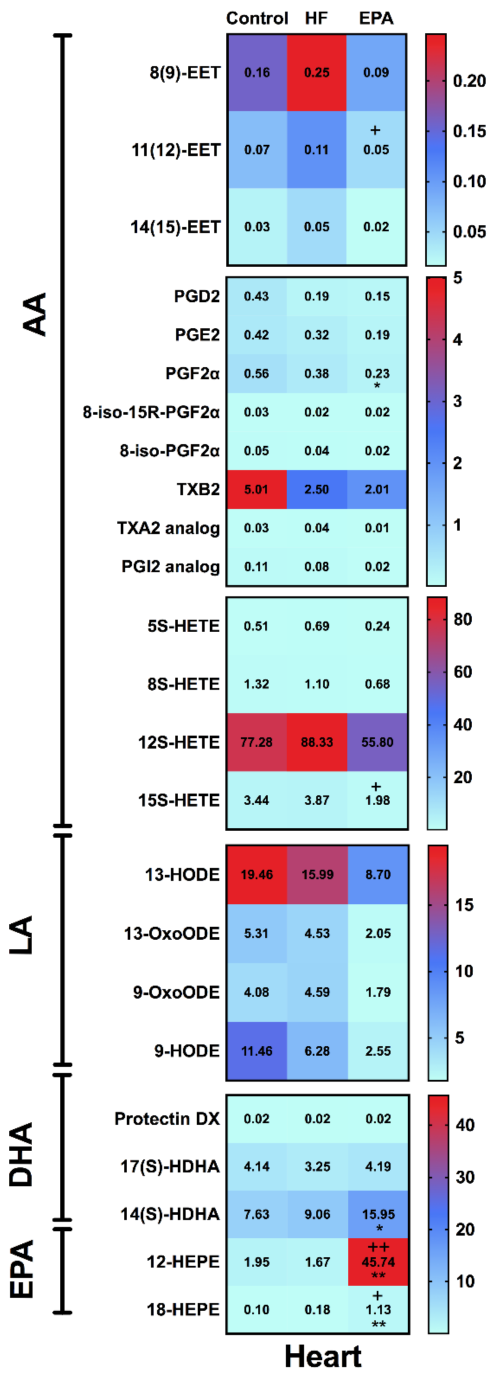


**Supplemental Figure 1. EPA ethyl esters increase levels of EPA derived lipid metabolites 12-HEPE and 18-HEPE.** Mass spectrometry based metabololipidomic analyses of cardiac tissue. Metabolites from eicosapentaenoic acid (EPA), docosahexaenoic acid (DHA), linoleic acid (LA) and arachidonic acid (AA) are depicted in the heat map. Male mice consumed experimental diets for 15 weeks. N=4-5 mice per diet. Data are average. *p<0.05, **p<0.01 as compared to a control diet and $^{+}$p<0.05, $^{++}$p<0.01 as compared to a high fat diet.

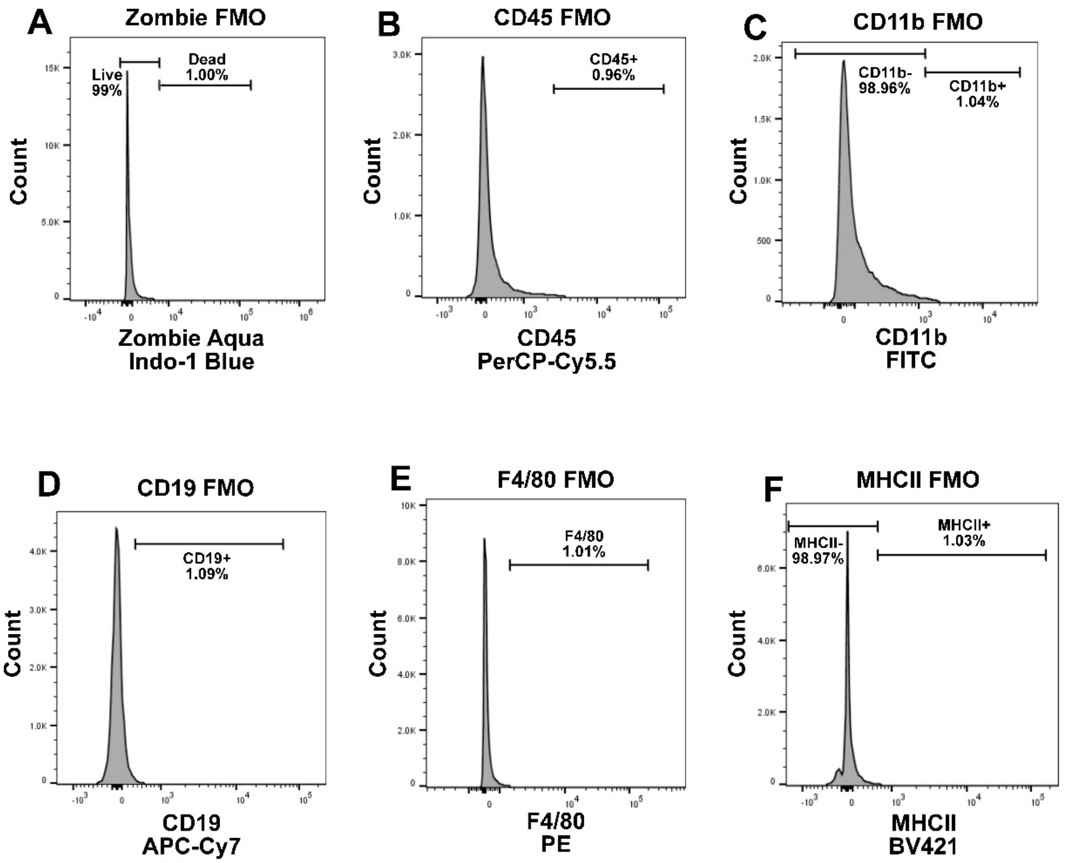


**Supplemental Figure 2.** Fluorescence Minus One gate for each fluorophore. The negative gates were drawn at a ~1% cutoff.

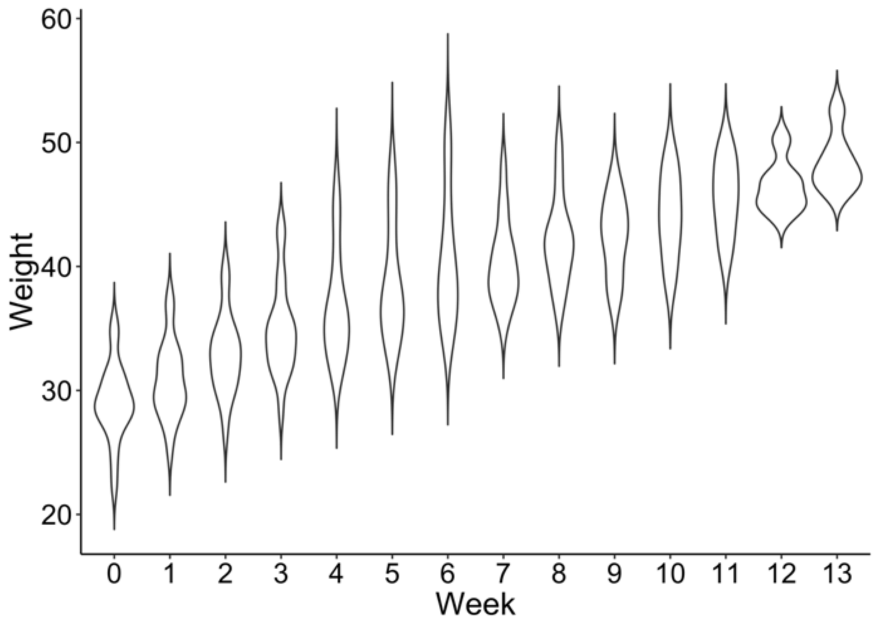


**Supplemental Figure 3. Body weights of DO mice consuming a high fat diet.** Violin plots of body weights as a function of time for DO mice at the completion of the first experimental design. This protocol entailed using DO mice that achieved approximately 14 grams of fat mass prior to RvE1 administration.
