## Supplemental Tables for "Resolvin E1 derived from eicosapentaenoic acid prevents hyperinsulinemia and hyperglycemia in a host genetic manner"

**Supplemental Table 1. Adipose metabolomics of C57BL/6J mice consuming a high fat diet in the absence or presence of EPA.**

| Full Metabolite Name | Shortened Metabolite Name | MetaboliteGroup | Type | EPA_containing | Log2FC | GreaterDiet | BH.adjust | EPA 122 | EPA 123 | EPA 125 | EPA 129 | EPA 130 | HF60 101 |
|---|---|---|---|---|---|---|---|---|---|---|---|---|---|
| Arg Lys Pro | Arg.Lys.Pro | Amino Acids | Amino Acids | NO | 2.391435934 | EPA | 0.004137 | 41894.47 | 38245.23 | 56102.32 | 58578.34 | 43989.35 | 8663.335 |
| Bouillonamide A | Bouillonamide.A | Macrolide | Macrolide | NO | 1.711871457 | EPA | 0.007652 | 2001.916 | 3728.059 | 2887.93 | 3233.896 | 4153.081 | 1078.604 |
| C-12 NBD-dihydro-Ceramide | C-12.NBD.dihydro.Ceramide | NBD.dihydro.Ceramide | Ceramide | NO | 4.485092 | EPA | 0.016708 | 672955 | 542882.1 | 654727.6 | 730481.1 | 478854.4 | 27675.25 |
| DG(20:5(5Z,8Z,11Z,14Z,17Z)/ | DG(40:10) | Diacylglycerols | DG | YES | 4.559313244 | EPA | 0.016708 | 1547479 | 1255585 | 1513081 | 1675377 | 1083498 | 57893.07 |
| DG(20:5(5Z,8Z,11Z,14Z,17Z)/ | DG(42:10) | Diacylglycerols | DG | YES | 1.738211816 | EPA | 0.005847 | 292851.9 | 226247.3 | 289155.9 | 329190.8 | 190577.6 | 75241.01 |
| DG(20:5(5Z,8Z,11Z,14Z,17Z)/ | DG(42:11) | Diacylglycerols | DG | YES | 2.118246586 | EPA | 0.004137 | 324616.8 | 266936.5 | 309371.6 | 359380.9 | 207733.8 | 62216.59 |
| DG(18:3(9Z,12Z,15Z)/20:4(5 | DG(38:7) | Diacylglycerols | DG | NO | 5.106853826 | EPA | 0.016708 | 591422.6 | 422151.2 | 628096 | 655187.3 | 360216.1 | 13433.45 |
| GalCer(d18:1/24:0) | GalCer(d42:1) | Ceramide | Ceramide | NO | 1.529804365 | EPA | 0.023611 | 8047.386 | 7886.809 | 12235.32 | 11425.62 | 8383.121 | 3561.868 |
| Gln His Thr | Gln.His.Thr | Amino Acids | Amino Acids | NO | 2.576216408 | EPA | 0.016708 | 4723.736 | 2775.599 | 7812.237 | 7943.821 | 16620.54 | 701.1674 |
| Gln Pro Pro | Gln.Pro.Pro | Amino Acids | Amino Acids | NO | 5.075716094 | EPA | 0.010623 | 19884.19 | 15018.81 | 15175.21 | 25062.05 | 34061.88 | 601.2257 |
| LysoPE(22:5(7Z,10Z,13Z,16Z | LysoPE(22:5) | Phosphoethanolamines | LysoPE | NO | 1.764144874 | EPA | 0.007334 | 1982.953 | 2203.75 | 3399.473 | 2836.308 | 2298.321 | 898.3778 |
| MGDG(18:2(9Z,12Z)/18:2(9Z | MGDG(36:4) | Galactolipid | Galactolipid | NO | 1.687142212 | EPA | 0.004137 | 22238.4 | 19629.72 | 28516.85 | 32115.17 | 26479.77 | 8665.813 |
| PA(12:0/17:0) | PA(29:0) | Phosphatidic Acids | PA | NO | 2.937431556 | EPA | 0.007652 | 15879.7 | 14858.32 | 19699.08 | 25203.4 | 11927.55 | 2266.021 |
| PA(18:3(9Z,12Z,15Z)/17:0) | PA(35:3) | Phosphatidic Acids | PA | NO | 3.255724591 | EPA | 0.016708 | 42002.08 | 28932.78 | 34168.86 | 41965.23 | 22321.74 | 3606.446 |
| PA(17:0/22:6(4Z,7Z,10Z,13Z | PA(39:6) | Phosphatidic Acids | PA | NO | 3.785281402 | EPA | 0.016708 | 7638.216 | 5490.414 | 7429.427 | 7544.938 | 4383.755 | 208.382 |
| PA(20:0/22:6(4Z,7Z,10Z,13Z | PA(42:6) | Phosphatidic Acids | PA | NO | 2.420649931 | EPA | 0.016708 | 74219.69 | 63436.55 | 99286.06 | 98334.77 | 75512.99 | 15055.03 |
| PC(18:3(6Z,9Z,12Z)/20:5(5Z, | PC(38:8) | Phosphocholines | PC | YES | 2.722774527 | EPA | 0.002979 | 4101.255 | 5884.606 | 4602.603 | 5707.123 | 6459.831 | 930.0704 |
| PC(16:1(9Z)/20:4(5Z,8Z,11Z, | PC(36:5) | Phosphocholines | PC | NO | 2.472924133 | EPA | 0.016708 | 10474.66 | 15040.6 | 11436.22 | 15269.43 | 17119.56 | 2475.902 |
| PC(22:4(7Z,10Z,13Z,16Z)/15 | PC(37:5) | Phosphocholines | PC | NO | -2.900490407 | HF | 0.016708 | 2977.967 | 1935.826 | 3053.077 | 2062.363 | 2362.217 | 20396.75 |
| PC(17:2(9Z,12Z)/22:6(4Z,7Z, | PC(39:8) | Phosphocholines | PC | NO | -2.282567229 | HF | 0.016708 | 2802.963 | 2917.057 | 3138.978 | 4667.757 | 2869.894 | 17602.83 |
| PE(20:5(5Z,8Z,11Z,14Z,17Z)/ | PE(36:5) | Phosphoethanolamines | PE | NO | 4.467561729 | EPA | 0.009449 | 50903.57 | 71080.22 | 58075.29 | 92887.58 | 115453.3 | 3197.142 |

Full metabolite names, shortened metabolite names, the metabolite group and type, EPA (22:5) containing metabolites (labeled as YES/NO), Log 2 fold changes (EPA/HF), the diet with the higher abundance of the metabolite (GreaterDiet), Benjamini-Hochberg adjusted p-values, and the raw metabolomic data are listed for all adipose metabolites below a BH p-value of 0.05 and Log2FC +/- 1.5. Each metabolite was tested for normality, followed by a T-test or Wilcoxon Rank-Sum Test and a BH-post hoc correction.

**Supplemental Table 2. Liver metabolomics of C57BL/6J mice consuming a high fat diet in the absence or presence of EPA.**

| Full Metabolite Name | Shortened Metabolite Name | MetaboliteGroup | Type | EPA_containing | Log2FC | GreaterDiet | BH.adjust | EPA 122 | EPA 123 | EPA 125 | EPA 129 | EPA 130 | HF60 101 | HF60 106 | HF60 108 | HF60 110 |
|---|---|---|---|---|---|---|---|---|---|---|---|---|---|---|---|---|
| Anandamide (20:5, n-3) | AEA(20:5, n-3) | Anandamide (AEA) | Anandamide | YES | 1.77397947 | EPA | 0.013388 | 5544.752 | 9706.924 | 11790.29 | 7592.302 | 9388.817 | 2357.634 | 2535.948 | 1960.771 | 3443.565 |
| Arg Asp Lys | Arg.Asp.Lys | Amino Acids | Amino Acids | NO | 1.690842337 | EPA | 0.03356 | 4620.197 | 19906.6 | 9057.827 | 11523.84 | 13019.46 | 1677.761 | 5617.889 | 2838.67 | 4269.591 |
| C-12 NBD-dihydro-Ceramide | C.12.NBD.dihydro.Ceramide | Ceramide | Ceramide | NO | 2.451081568 | EPA | 0.007747 | 10463.06 | 16019.61 | 9227.833 | 13892.19 | 13687.45 | 3279.367 | 2155.511 | 2348.021 | 1476.377 |
| Cer(d18:1/16:0) | Cer(d34:1) | Ceramide | Ceramide | NO | 2.806386276 | EPA | 0.019909 | 258567.7 | 24724.76 | 18302.2 | 46148.13 | 25671.25 | 11874.42 | 12609.47 | 7067.425 | 11153.23 |
| Chivosazole A | Chivosazole.A | Macrolide | Macrolide | NO | 2.230318498 | EPA | 0.019909 | 5544.418 | 23564.94 | 15462.14 | 19384.21 | 25806.14 | 4561.172 | 3281.618 | 4022.359 | 3438.289 |
| DG(20:5(5Z,8Z,11Z,14Z,17Z)/ | DG(40:10) | Diacylglycerols | DG | YES | 2.46414865 | EPA | 0.007747 | 23530.11 | 35746.75 | 19845.19 | 29930.08 | 28854.08 | 7229.963 | 4803.678 | 4693.606 | 3266.36 |
| DG(20:5(5Z,8Z,11Z,14Z,17Z)/ | DG(42:11) | Diacylglycerols | DG | YES | 2.039717555 | EPA | 0.019909 | 11097.93 | 3923.338 | 2991.482 | 2748.868 | 3084.674 | 1937.356 | 970.5071 | 1147.446 | 584.4419 |
| DG(18:3(9Z,12Z,15Z)/20:4(5Z | DG(38:7) | Diacylglycerols | DG | NO | -1.454841989 | HF | 0.045162 | 124.2307 | 2269.188 | 2845.917 | 1539.615 | 1970.302 | 6350.403 | 2968.094 | 6651.087 | 3217.633 |
| DG(20:1(11Z)/20:3(8Z,11Z,14 | DG(40:4) | Diacylglycerols | DG | NO | -1.857840044 | HF | 0.035631 | 414.5454 | 1591.887 | 3075.688 | 432.7505 | 977.9517 | 6765.352 | 3022.686 | 6120.513 | 2918.794 |
| Docosahexaenoic Acid | DHA | DHA | DHA | NO | 2.093197428 | EPA | 0.012841 | 6307.439 | 11071.47 | 14491.03 | 8609.298 | 10978.73 | 2129.609 | 2069.175 | 1364.805 | 4084.19 |
| 3b,16a-Dihydroxyandrostenc | Dihydroxyandrostenone.sulfate | Sulfated steroids | Sulfated Steroids | NO | 1.96847808 | EPA | 0.013388 | 11954.14 | 27967.65 | 27321.35 | 18912.04 | 22915.02 | 3666.087 | 6275.025 | 3135.668 | 9219.126 |
| 5,7-Dihydroxy-4'-methoxy-8- | flavanone | Flavanone | Flavanone | NO | 1.773772164 | EPA | 0.019909 | 18642.88 | 6532.969 | 21835.32 | 13081.24 | 19067.81 | 4479.47 | 3854.67 | 2387.179 | 7798.568 |
| alNAcbeta1-4(NeuGcalpha2-3 | Galactosylceramide(40:1) | Ceramide | Ceramide | NO | -2.478892208 | HF | 0.019909 | 63.55114 | 8998.725 | 6000.511 | 3711.413 | 8571.961 | 35945.59 | 22819.03 | 41036.7 | 22155.83 |
| alNAcbeta1-4(NeuGcalpha2-3 | Galactosylceramide(42:2) | Ceramide | Ceramide | NO | -2.648712069 | HF | 0.019909 | 51.88366 | 2932.832 | 2233.087 | 1409.415 | 2700.981 | 11711.04 | 7210.651 | 15555.01 | 12321.55 |
| GalCer(d18:1/23:0) | GalCer(d41:1) | Ceramide | Ceramide | NO | -2.269075549 | HF | 0.007747 | 1373.163 | 15020.28 | 12032.07 | 12290.28 | 12843.27 | 55005.07 | 38233.4 | 58684.57 | 54606.76 |
| GalCer(d18:1/24:0) | GalCer(d42:1) | Ceramide | Ceramide | NO | -2.51113696 | HF | 0.012681 | 1244.782 | 19636.7 | 13633.65 | 10585.63 | 10700.94 | 70605.63 | 53197.37 | 79620.25 | 51063.36 |
| Gln His Thr | Gln.His.Thr | Amino Acids | Amino Acids | NO | 3.198828512 | EPA | 0.019909 | 14619.02 | 56504.84 | 40374.21 | 31303.57 | 39140.37 | 2275.044 | 4568.387 | 2306.273 | 6702.135 |
| Gln Pro Pro | Gln.Pro.Pro | Amino Acids | Amino Acids | NO | 5.488421468 | EPA | 0.01494 | 39666.98 | 74021.31 | 33478.31 | 46037.73 | 36157.77 | 1011.432 | 871.2841 | 802.8488 | 1401.694 |
| Ivermectin B1b | Ivermectin.B1b | Lactone | Lactone | NO | 1.602176418 | EPA | 0.024292 | 1305.993 | 4252.53 | 4142.477 | 5564.524 | 5344.6 | 1.00E-05 | 1951.182 | 1296.642 | 2183.021 |
| LysoPE(20:5(5Z,8Z,11Z,14Z,1 | LysoPE(20:5) | Phosphoethanolamines | LysoPE | YES | 3.279364063 | EPA | 0.019909 | 4486.95 | 26065.36 | 13399.95 | 14635.8 | 17630.4 | 2847.858 | 389.3203 | 2752 | 290.8725 |
| Macaflavone II | Macaflavone.II | Flavonoids | Flavanone | NO | 1.487277655 | EPA | 0.000994 | 7015.207 | 7118.376 | 8670.256 | 6795.163 | 7589.154 | 3165.605 | 2474.736 | 1896.907 | 3074.318 |
| Methylsyringin | Methylsyringin | Glucoside | Glucoside | NO | 1.510848086 | EPA | 0.019909 | 3731.94 | 5056.865 | 3272.364 | 4056.697 | 4533.18 | 860.1353 | 1523.693 | 304.9037 | 3108.51 |
| (2R,6x)-7-Methyl-3-methylen | octanetetrol.2.glucoside. | Glucoside | Glucoside | NO | 1.983520777 | EPA | 0.012681 | 54919.93 | 89409.12 | 110605.6 | 71151.7 | 84976.09 | 18161.23 | 19857.25 | 16118.41 | 29020.05 |
| Oxo-Ergotamine | Oxo.Ergotamine | Ergopeptines | Ergopeptines | NO | 4.437344148 | EPA | 0.019909 | 519.9044 | 5112.722 | 3086.143 | 2678.54 | 3973.033 | 62.25989 | 177.7969 | 71.35096 | 256.1367 |

Full metabolite names, shortened metabolite names, the metabolite group and type, EPA (22:5) containing metabolites (labeled as YES/NO), Log 2 fold changes (EPA/HF), the diet with the higher abundance of the metabolite (GreaterDiet), Benjamini-Hochberg adjusted p-values, and the raw metabolomic data are listed for all liver metabolites below a BH p-value of 0.05 and Log2FC +/- 1.5. Each metabolite was tested for normality, followed by a T-test or Wilcoxon Rank-Sum Test and a BH-post hoc correction.

**Supplemental Table 3. Single Nucleotide Polymorphisms in EPA/RvE1 metabolizing genes.**

| Variant name | chr | chr.start | chr.end | Minor allele | Global minor allele frequency | Gene |
|---|---|---|---|---|---|---|
| rs1046587 | 14 | 24316854 | 24316854 | A | 0.2716 | BLT1 |
| rs1046584 | 14 | 24316578 | 24316578 | T | 0.2358 | BLT1 |
| rs2224123 | 14 | 24313752 | 24313752 | T | 0.1567 | BLT1 |
| rs4981503 | 14 | 24317087 | 24317087 | T | 0.155 | BLT1 |
| rs3181384 | 14 | 24317770 | 24317770 | T | 0.1532 | BLT1 |
| rs2224122 | 14 | 24314208 | 24314208 | C | 0.111 | BLT1 |
| rs3742510 | 14 | 24314475 | 24314475 | C | 0.07188 | BLT1 |
| rs3742511 | 14 | 24315705 | 24315705 | C | 0.07169 | BLT1 |
| rs551001577 | 10 | 45407402 | 45407402 | C | 0.05092 | 5LOX |
| rs17157756 | 10 | 45385220 | 45385220 | G | 0.05112 | 5LOX |
| rs115074014 | 10 | 45385210 | 45385210 | A | 0.05112 | 5LOX |
| rs12266963 | 10 | 45433456 | 45433456 | G | 0.05471 | 5LOX |
| rs71494783 | 10 | 45432341 | 45432341 | G | 0.05471 | 5LOX |
| rs115200794 | 10 | 45427617 | 45427617 | C | 0.05671 | 5LOX |
| rs72796464 | 10 | 45410446 | 45410446 | C | 0.05851 | 5LOX |
| rs55661611 | 10 | 45424378 | 45424378 | T | 0.0607 | 5LOX |
| rs7080977 | 10 | 45421868 | 45421868 | C | 0.0615 | 5LOX |
| rs12783037 | 10 | 45434517 | 45434517 | C | 0.0617 | 5LOX |
| rs11239501 | 10 | 45383352 | 45383352 | A | 0.0619 | 5LOX |
| rs7077173 | 10 | 45421751 | 45421751 | G | 0.0621 | 5LOX |
| rs17444064 | 10 | 45396252 | 45396252 | C | 0.0625 | 5LOX |
| rs41526545 | 10 | 45395767 | 45395767 | G | 0.0637 | 5LOX |
| rs55700434 | 10 | 45391508 | 45391508 | T | 0.0637 | 5LOX |

SNPs in genes of the EPA/RvE1 pathway are listed by reference SNP (RS) number, chromosome number, chromosome locations (chr.start and chr.end), minor allele, global minor allele frequency, and the gene the SNP is contained within.
